# Optimizing Cancer Vaccinations Using a Physiologically Based Pharmacokinetic (PBPK) Model

**DOI:** 10.1101/2025.11.17.688910

**Authors:** Mohammad R. Nikmaneshi, Timothy P. Padera, Lance L. Munn

**Affiliations:** Department of Radiation Oncology, Massachusetts General Hospital and Harvard Medical School, Boston, MA 02114, USA

## Abstract

Antigen-based tumor vaccines rely on adjuvants to stimulate local inflammation, recruit antigen-presenting cells (APCs), and enhance immune activation. However, the complex interplay between antigen transport, lymphatic drainage, and immune cell dynamics across organs remains poorly understood, limiting the rational design of vaccination strategies. Here, we present a multiscale compartmental Physiologically Based Pharmacokinetic (PBPK) model of antigen vaccination that integrates systemic circulation, lymphatic connectivity, and immune cell activation at the whole-body level. The model incorporates arterial, venous, and lymphatic flows, organ-specific interstitium and lymph node (LN) networks, and a superficial skin network. The model reproduces spatiotemporal distributions of antigen and suppressive factors, APC activation, and nT priming across activation sites, including LNs and spleen. Our results show that the sensitivity of vaccination-induced immunity is highly related to antigen and suppressive factor production by the tumor, and that early-stage vaccination, enhances immunity. Since the model is able to identify optimal vaccination administration over the course of tumor growth for each patient with certain levels of antigen and immune suppressive factors, it can serve as the foundation for digital twins of patients to help inform anti-cancer vaccination strategies.

**Teaser:** A PBPK model links body-wide immune transport to optimize cancer vaccine design and delivery.

## Introduction

Despite encouraging results from cancer vaccines—particularly mRNA-based and antigen-based platforms—patient responses remain highly variable, even among individuals with the same tumor type and stage ^1-4^. Preclinical and animal studies have consistently demonstrated considerable fluctuations in response to immunotherapy, even within genetically identical models ^3,5^. One of the main reasons for this inconsistency is the complex and heterogeneous trafficking of T cells to the tumor and their variable infiltration patterns, an aspect captured by the concept of tumor-immune “hotness,” which refers to the quality and spatial distribution of T cell infiltration within the tumor microenvironment (TME). Tumors are generally classified immunologically into three categories: immune-desert or “cold” tumors characterized by absent or non-activated T cells; immune-excluded or non-inflamed “cold” tumors in which T cells are present but fail to infiltrate the tumor parenchyma; and immune-inflamed or “hot” tumors that are enriched with infiltrating, activated T cells ^6-10^. The effectiveness of cancer vaccination can therefore be strongly influenced by the tumor’s hotness.

In addition to the tumor properties, three major factors —the vaccination site, timing of administration, and type of vaccination— may play critical roles in determining the efficacy of cancer vaccination. As a tumor grows, the distributions of immune agents such as antigen-presenting cells (APCs), antigens, naïve T cells, and cytokines undergo dynamic changes throughout the body ^11,12^. Consequently, the potential for T cell activation, which depends on pro- and anti-inflammatory signals, also varies with time and location in the body. Understanding the spatiotemporal distributions of these components is crucial to the success of any immunization strategy, including cancer vaccines.

Beyond local immune evasion at the tumor site, tumors can exert systemic immunosuppressive effects by releasing soluble factors that interfere with immune priming in secondary lymphoid organs. For example, tumor-derived cytokines and metabolites such as TGF-β, IL-10, VEGF, and lactic acid have been shown to suppress dendritic cell (DC) maturation and function, impairing antigen presentation and co-stimulation required for naive T cell activation in lymph nodes ^13-17^. These factors can downregulate MHC and costimulatory molecules (e.g., CD80, CD86) on APCs, inhibit their migration to lymph nodes, and promote the development of tolerogenic DCs. In parallel, they can directly inhibit T cell receptor signaling, reduce IL-2 production, and promote anergy or regulatory phenotypes in naive T cells, even before they reach the tumor ^12,18-20^. This pre-conditioning of the systemic immune environment compromises the initiation of effective anti-tumor responses and may contribute to poor responsiveness to immunotherapies.

Common routes of vaccine administration, such as pulmonary delivery via inhalation, oral or gastrointestinal (GI) delivery, subcutaneous injection, and intramuscular injection, each engage the immune system differently depending on the tumor’s location, stage, and systemic immunosuppressive activity ^21-25^. Key physiological and anatomical variables—including blood and lymphatic circulation, tissue-specific architecture, lymphatic connectivity, and tissue transport properties—strongly affect the distribution of immune components and thus the resulting vaccine performance ^26-30^. These complexities mean that cancer vaccination outcomes can be highly variable and may differ significantly not only between species but also among individual patients. Due to the anatomical and physiological differences between humans and animals, preclinical animal models often fail to recapitulate human vaccination responses accurately ^31-34^. Furthermore, there is currently no clinically available tool to optimize vaccination strategies for individual cancer patients.

In this context, computational modeling emerges as a powerful and efficient approach to predict patient-specific immune responses to vaccination, with minimal time investment and high biological relevance. Although several mathematical models have been developed to simulate the systemic trafficking of immune cells and the local immune dynamics in the tumor microenvironment ^35-40^, a comprehensive whole-body model that incorporates both systemic immunity and cancer metabolism remains unavailable. Such a model is critical for generating a holistic understanding of the immune landscape and for predicting how the body will respond to various vaccination strategies—including variations in timing, site, and vaccine type—while accounting for the tumor’s biological behavior and stage.

The study of systemic immune responses to immunization requires comprehensive models that capture the complexity of immune dynamics throughout the entire body. Recently, Physiologically-based pharmacokinetic/pharmacodynamic (PBPK/PD) models and multi-compartmental frameworks have been developed to map the immune system’s behavior across organs, incorporating vascular and lymphatic connectivity ^37,41-43^. These models are instrumental in addressing critical questions about the timing, magnitude, and localization of immune responses to various cancer therapies, including vaccination, checkpoint blockade, adoptive T cell therapy such as CAR-T, and radiotherapy ^37,44^. Some early models simplify systemic immunity by including only a few compartments—typically the target organ (e.g., tumor), the bloodstream, and a single lymph node—serving as the site for T cell priming ^45-48^. While useful for initial insights, these models often fail to capture the heterogeneity of immune cell trafficking, recirculation, and clearance mechanisms that can delay or dampen immune activation in patients. More advanced approaches incorporate multiple lymphoid and non-lymphoid compartments to better resolve immune cell kinetics, account for physiological clearance, and improve prediction of treatment outcomes^37,49,50^. Recent innovations in systems immunology have aimed to bridge these gaps by developing integrated models that simulate the spatiotemporal dynamics of immune responses at systemic levels. These include models that account for organ-specific lymphatic drainage, allowing simultaneous tracking of T cell activation across multiple lymph nodes and their migration toward tumors ^37,51^. Such frameworks are particularly valuable for studying responses to combination therapies, like radiotherapy (RT) and vaccination, which can initiate immune activation in distinct anatomical contexts ^52,53^. Additionally, the heterogeneity of organ-specific environments—ranging from mechanical properties and vascular permeability to cytokine gradients and antigen presentation efficiency—adds further complexity to accurately predicting immune responses ^54-60^. Simplified models often overlook these factors, leading to poor translational relevance in predicting post-vaccination immunity or T cell–based therapy outcomes. Therefore, incorporating organ-specific immunobiology, lymphatic topology, and dynamic inter-compartmental feedback is essential for building predictive and clinically relevant models of immunotherapy. Another issue with current PBPK models is their lack of species specificity. It is often unclear whether the model has been developed for humans, mice, or another species. Since each species has distinct anatomical and physiological characteristics, using a single model for both humans and mice can lead to irrelevant results and poor translational outcomes for patients.

To address these gaps, we have developed a computational systems immunology model of human immuno-physiology using a compartmental PKPD approach that integrates the circulatory and lymphatic systems, major internal organs, the superficial immune network of the skin, lymph node architecture, immune cell trafficking and tumor metabolism (Figure 1). The model uses a matrix-based spatial framework that allows dynamic configuration of interconnected compartments in the human body. Each compartment includes vascular and tissue-level structures responsible for interstitial fluid flow and molecular transport and is linked to lymph nodes either in parallel or in series, capturing physiologically accurate transport pathways. The skin is modeled as a special organ, and includes multiple interconnected subcompartments, each with draining lymph nodes that are also integrated into the systemic network, thereby linking peripheral immune responses to systemic immunity. This whole-body framework serves as a virtual platform for simulating and optimizing cancer vaccination strategies in a patient-specific manner across spatial and temporal scales.

**Figure 1.**
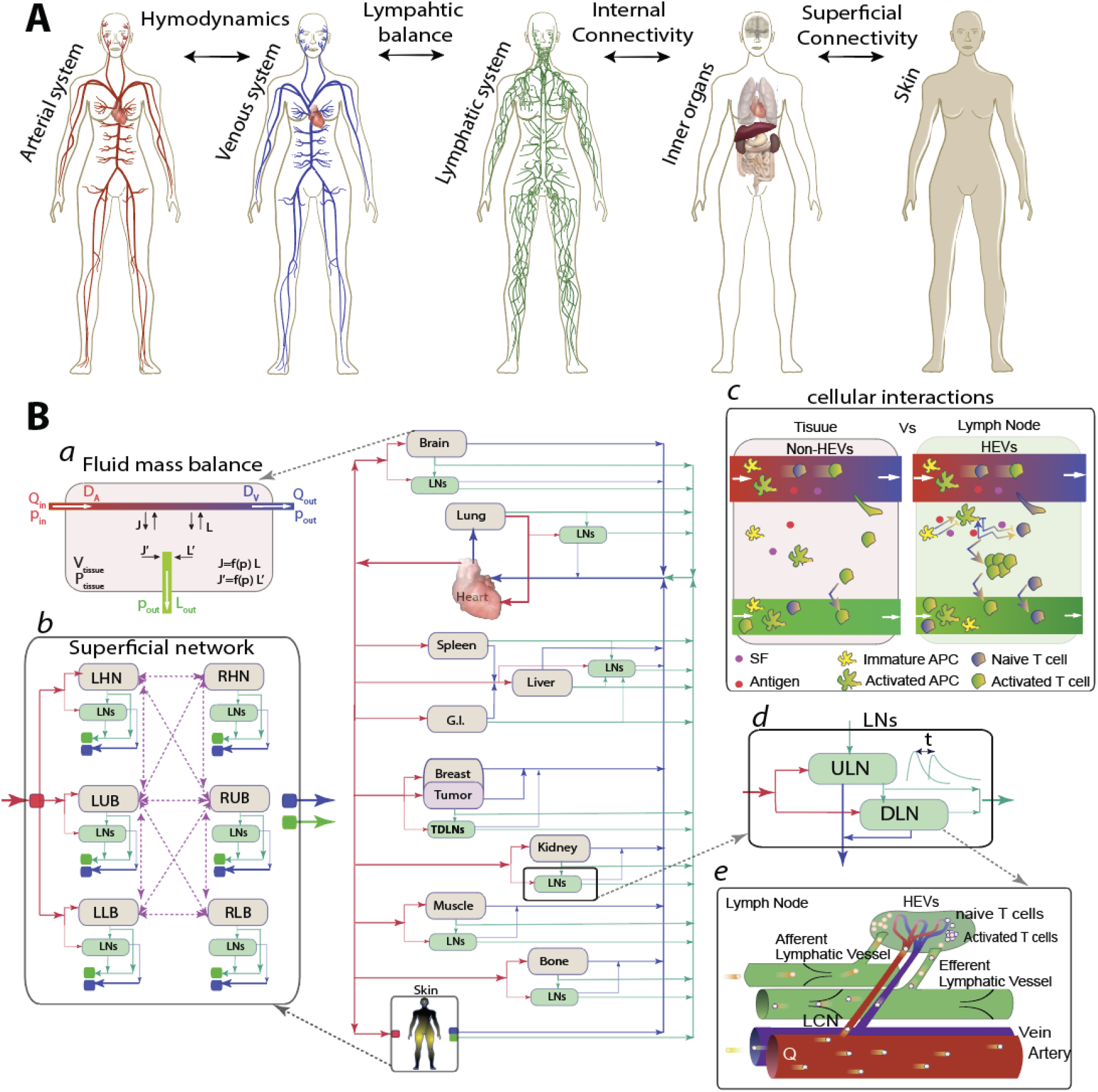
Model configuration and computational PKPD model. (A) The model integrates multiple body layers, including the arterial, venous, and lymphatic systems, internal organs, and skin. (B) The computational diagram illustrates how individual internal organs, represented as separate compartments, are connected via arterial (red arrows) and venous (blue arrows) circulations, as well as lymphatic drainage (green arrows). Each compartment has its own fluid mass balance through blood vessels, interstitium, and the lymphatic system (*a*). Skin has its own superficial network including different lymphosomes interconnected with superficial flows (dashed-purple arrows) and also interconnected with other inner organs through blood and lymphatic local recirculation (*b*). signaling and localization of APCs and T cells under antigen and SF is different in interstitium of tissue and LNs (*c*). Each compartment contains its own LN network, consisting of ULNs and DLNs (*d*). Each LN includes all physiological pathways: afferent and efferent lymphatic vessels, and blood circulation through high endothelial venules (HEVs) (*e*).

## Methods

### Scales and agents of the model

The components of the compartmental PKPD model are illustrated in Fig.1. The model is multiscale, including body-scale, cellular-scale, and molecular-scale.

At the ***body scale***, the model incorporates four layers: arterial, venous, and lymphatic circulations, and organs, which include the internal organs and superficial (skin) tissue (Fig.1A). The major internal organs include the heart (with distinct left and right ventricles), brain, lungs, liver, spleen, GI tract, kidneys, breast tissue, skeletal muscle, and bone. The superficial skin network is divided into six regions: left and right head-neck (LHN and RHN), left and right upper body (LUB and RUB), and left and right lower body (LLB and RLB). The tumor compartment can be embedded in any organ, including the skin. The computational chart shows inner organs and skin compartments with their associated draining LNs interconnected by arterial (red arrows), venous (blue arrows), lymphatic (green arrows), and superficial, interstitial (purple dashed arrows) flows (Fig.1B). Each compartment includes tissue, blood vessels and lymphatic vessels (Fig.1B.a).

Using hemodynamic parameters of each compartment—including blood flow rate (Q), luminal pressure, vessel wall permeability, and porosity—the arterial and venous flows are coupled with lymphatic flows throughout the system. A modified Starling’s law with interstitial fluid pressure and oncotic pressure determined by vessel wall permeability/porosity defines the lymphatic balance (Fig.1B.*a*). The coupled blood–lymphatic system provides internal connectivity among organs and superficial connectivity across skin compartments, which in turn link to inner organs (Fig.1B.*b*). Each organ receives physiological blood flow based on known flow rates, and organ-specific transport properties regulate exchanges of cells and molecules between the vasculature and interstitium. This framework allows the model to simulate dynamic infiltration and recirculation of immune components between systemic compartments and target tissues, providing a platform to explore vaccination strategies and immune responses across the whole body. Additional details are provided in the Supplementary Materials.

At the ***cellular scale***, the model includes tumor cells and angiogenic vasculature, naive T cells (nTs), effector T cells (eTs), immature antigen presenting cells (imAPCs) and activated antigen presenting cells (aAPCs) (Fig.1B.*c*). CD4/CD8 post-activation ratios are assumed constant among eTs. nTs are generated in the thymus, modeled as a variable source with cytokines directly injected into the left ventricle. Generation of imAPCs mainly occurs in bone marrow and, to a lesser extent, in skin, and is affected by cytokine levels.

At the ***molecular scale***, the model simulates transport and interactions of antigens, cytokines, VEGF, IL-2, and a putative tumor-derived suppressive factor (SF). Antigen is released by cancer cells or supplied via vaccination; inflammatory cytokines are considered as a single species, which is secreted strongly by aAPCs and moderately by imAPCs and other immune cells (assumed a constant source for other immune cells here). VEGF is released by tumor cells to promote angiogenesis; IL-2 is considered separately, and is secreted by activated eTs (both CD4 and CD8); SF is released by cancer cells.

### Vaccine model

We introduce the vaccine as a pulse function of antigen source within the target tissue. Since the antigen vaccines are usually infused with adjuvant to induce inflammation and stimulate APC generation and recruitment, we assume that, in addition to the antigen source, the vaccine includes adjuvant. Thus, the model accurately simulates the complete loop of vaccination-induced APC–cytokine dynamics: vaccination releases adjuvant at the injection site, which enters the systemic circulation, enhances APC generation and recruitment, leading to cytokine release by APCs, which in turn further amplify APC responses.

### Inhibition by tumor-produced suppressive factors

Since cancer can release many types of metabolites, including suppressive factors (SFs) that inhibit T cell activation, understanding their role in modulating immune response is essential. Although the precise mechanisms by which SFs influence T cell priming remain unclear, we assume in this model that they affect both the activation of antigen-presenting cells (APCs) and the ability of the activated APCs to activate naïve T cells (nTs). Therefore, in addition to tracking the distributions of antigens, nTs, and APCs within lymph nodes (LNs), the distribution of SFs within the LN network is considered a critical parameter in determining the extent of systemic T cell activation.

### Cellular dynamics

#### Combining cellular and molecular scales

imAPCs and aAPCs, nTs, and eTs interact differently within the tissue interstitium compared to the LN interstitium (Fig.1B/*c*). APCs can traffic to both tissue and LN interstitium. nTs can enter the LN interstitium but not the tissue interstitium ^61-63^. eTs can be generated and proliferate in the LN interstitium when nTs interact with aAPCs. Activation of nTs occurs when antigen level is sufficient, and SF level is low; eTs can exit the blood and migrate within tissue compartments, but do not enter LNs. eT cells can enter the tumor tissue and interact with cancer cells. As we are only interested in the initial activation process and the accumulation of T cells in the tumor, we neglect immunoregulatory mechanisms of T cells once they enter the tumor tissue, such as exhaustion. We also neglect later phases of immune response, such as memory cell formation and circulation. Key immune interactions, such as APC activation – which depends on the antigen, cytokine and SF levels – and subsequent nT activation, occur in the LN tissue, and not in other organs (Fig.1B.*c*). imAPCs circulate in the blood vasculature and home to compartments with high antigen level; once there, they can be activated, depending on the local levels of antigen and SF. aAPCs then migrate to the lymphatic circulation and can co-localize with nT cells in LNs to activate them; again, this step can be limited by SF.

#### Activation sites

in general, LNs and spleen are activation sites for both imAPCs and nTs. Each organ has multiple draining LNs with the potential for different antigen, cytokine and SF levels in each. We simplify the system by including two lymph nodes arranged in series that drain each organ, notated as the upstream LNs (ULNs) and downstream LNs (DLNs) (Fig.1B, *d*). Each LN has its own blood circulation, and cell trafficking through HEVs as well as afferent/efferent pathways of lymphatic circulation, are considered independently (Fig.1B, *e*).

## Results and discussion

To demonstrate the various systemic mechanisms involved in systemic T cell activation, we first simulated vaccination in the left arm of a patient with a tumor in the right breast (Figure 2). Since cancer can release many types of metabolites, including suppressive factors (SFs) that inhibit T cell activation, understanding their role in modulating immune response is essential. Although the precise mechanisms by which SFs influence T cell priming remain unclear, we assume in this model that they affect both the activation of antigen-presenting cells (APCs) and the ability of the activated APCs to activate naïve T cells (nTs). Therefore, in addition to tracking the distributions of antigens, nTs, and APCs within lymph nodes (LNs), the distribution of SFs within the LN network is considered a critical parameter in determining the extent of systemic T cell activation (Figure 2). Tumor growth is initiated on day 0 and progresses over 25 days. Antigen/adjuvant vaccination is administered subdermally one week after tumor initiation (i.e., when the tumor size is 15 mm^3^) and two weeks after vaccination. In this simulation, the patient is assumed to have a low level of tumor-derived antigen, which is negligible compared to the antigen introduced by the vaccine. Thus, immune activation is limited before vaccination. The results show the dynamics of key immune components in the ULNs and DLNs draining the vaccination site and tumor for two weeks post-vaccination, including antigen levels, SF levels, immature APCs (imAPCs), activated APCs (aAPCs), cytokines, naïve T cells (nTs), and effector T cells (eTs)(Figure 2). Vaccination administered on day 7 subdermally releases antigen into the local tissue, which subsequently drains into nearby LNs (both ULNs and DLNs). As the tumor grows, SFs are released into the systemic circulation, spreading to the vaccination site and its associated LNs. In response to the vaccine (antigen and adjuvant), APCs and other immune cells release cytokines that accumulate in the vaccination site and its draining LNs (Figure 2D).

**Figure 2.**
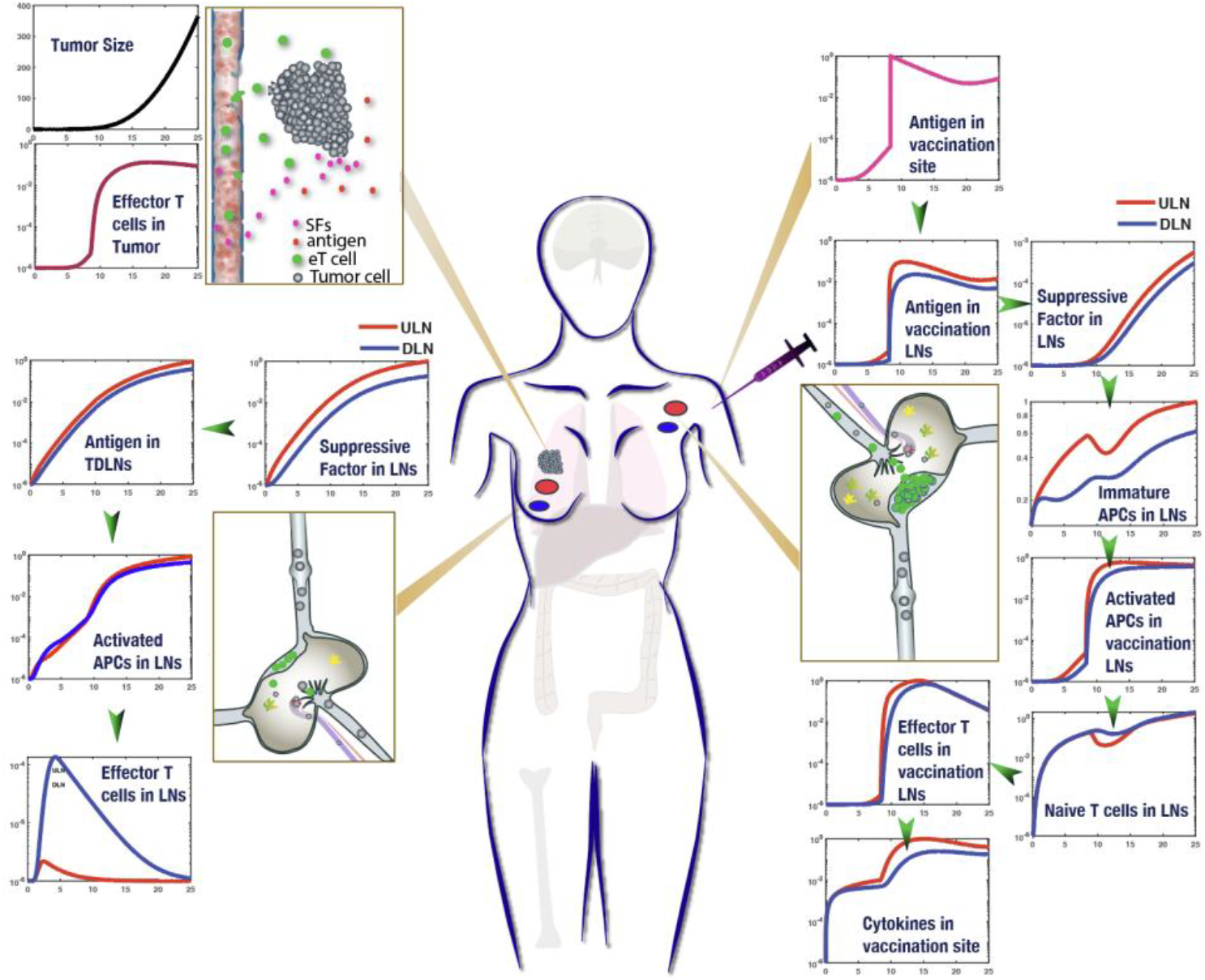
Mechanistic overview and temporal dynamics of tumor growth and immune response following vaccination. The plots show tumor size, antigen levels, suppressive factor (SF) levels, cytokine concentrations, immature and activated antigen-presenting cells (imAPCs and aAPCs), naïve T cells (nTs), and effector T cells (eTs). Tumor growth reflects the increase in tumor tissue volume over time. Antigen levels are normalized to the concentration in the vaccination site; SF levels are normalized to the concentration in the tumor tissue; cytokine levels are normalized to those in the vaccination site. imAPC and aAPC levels are normalized to the baseline imAPC level in the bone. nT and eT levels are normalized to the initial nT population released from the thymus into the heart. The model assumes that tumor-specific nTs are generated upon cancer initiation, and that the total nT population remains at steady-state during the simulation. Green arrows indicate the process pathway in the LNs.

Immature APCs, which are primarily generated in the bone marrow and to a lesser extent in the skin, enter the systemic circulation and migrate to peripheral tissues, where they can be activated depending on the local levels of antigen and SF. Naïve T cells, released from the thymus into the blood circulation, are distributed systemically and accumulate in LNs, where they can differentiate into effector T cells following stimulation by aAPCs; this step is enhanced by the presence of antigen and inhibited by SF. Because LNs are the primary sites for colocalization of APCs and T cells, the activated populations of APCs and T cells are typically higher in LNs than in peripheral tissues.

Because the suppressive factor can modulate activation of APCs and T cells, it is important to consider its spatiotemporal variation after vaccination at various sites. While the tumor grows at any SF rate, the spatiotemporal distribution of SF in the LNs and spleen dynamically changes, thereby influencing the activation rates of APCs and T cells. To simplify the presentation of the results, we define a parameter – the activation potential (AP) of a LN or spleen, which is related to the probability of activation occurring at that location. In the model, AP serves as a probabilistic reduction factor modulating APC and T-cell activation rates according to SF concentration (see Eqs. 10–13 in the Supplemental Material). We can then see how different SF production rates affect the AP for the ULNs, DLNs and spleen (Figure 3A). The AP is then integrated over the time course of the simulation (i.e., the area under the curve) to produce a cumulative AP, (Figure 3B, polar plot). This analysis helps identify the appropriate time and site for vaccination at each stage of the tumor growth. For low SF, tumor LNs have some activation potential when the vaccination is administered early (until day 5 for ULNs and until day 7 for DLNs). However, increasing SF by 40% shortens the effective vaccination times to day 2 and 5 and then non-activation and day 3 for ULNs and DLNs, respectively. Similar trends are seen for the spleen and LNs in other locations. In general, ULNs have lower activation potential, especially those draining the tumor, lung, G.I., muscle and superficial skin LNs. Interestingly, in the LNs draining the lungs, the ULN has considerably less activation potential than the DLNs at every SF production rate. This is due to the fact that all blood (and SF) passes through the lungs, and the SF gets concentrated in the ULNs faster than the DLNs.

**Figure 3.**
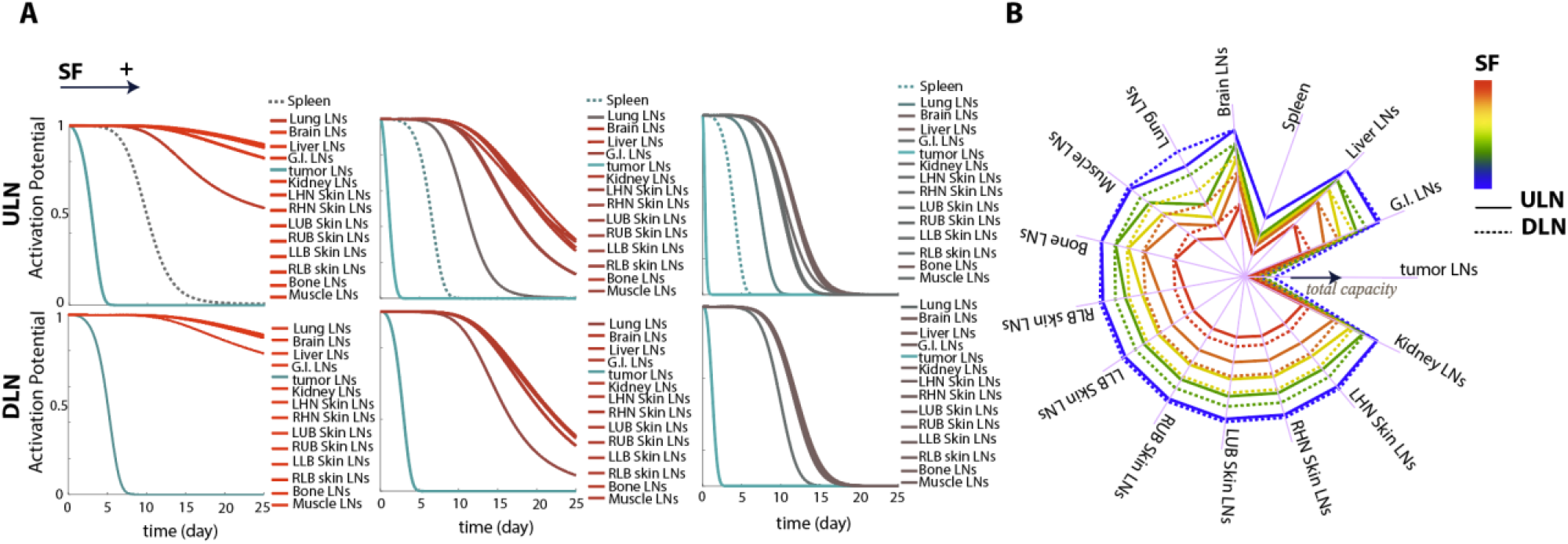
Activation potential of LNs and spleen. A) The plot colors transition from green to red, representing the activation potential of each site, ranging from non-active (green) to fully active (red), when vaccination begins on day 7. The first row corresponds to ULNs and the second row to DLNs. B) The polar plots display the area under the curve (AUC) of activation capacity profiles over 25 days of tumor growth. Solid lines represent ULNs, while dashed lines represent DLNs.

Post-vaccination T cell activation is determined by the co-localization of antigen, aAPCs, and naïve nTs within a LN, provided the LN is not suppressed by SF (Fig. 3). This is illustrated in the scenario shown in Figure 1.c: even when the tumor-draining LNs have high antigen level and contain both nTs and aAPCs, an elevated SF concentration can prevent T cell activation (maximum normalized activation of T cells in tumor-draining DLN 10^−4^ logscale vs 10^0^ logscale for the vaccination LNs). As a result, eTs are instead generated in other, less suppressed LNs and subsequently must traffic through the vasculature and migrate into the tumor tissue (Figure 1).

To investigate how the vaccination site affects the systemic immune response, we introduced vaccine at various internal organs and skin regions and monitored the distribution of eT cells in all LNs and the spleen. We simulated responses in patients with breast tumors exhibiting varying levels of antigen and SF, including: (1) low antigen and low SF, (2) high antigen and low SF, (3) low antigen and high SF, and (4) high antigen and high SF.

To assess the efficiency of vaccination, we define three metrics: i) activation site occupancy (ASO) calculated as the area under the curve (AUC) of eT cell concentration, calculated for the spleen and each LN during the two weeks following vaccination (Fig. 4), ii) Overall Immune Hotness (OIH), which is the total number of eTs present, integrated over all LNs and the spleen (Fig.5), and iii) Tumor Immune Hotness (TIH), which is the fraction of all eT cells that are localized in the tumor (Fig.6). Thus, ASO quantifies activation at each activation site, OIH is related to systemic immune activation, and TH is related to anti-tumor cytotoxicity.

**Figure 4.**
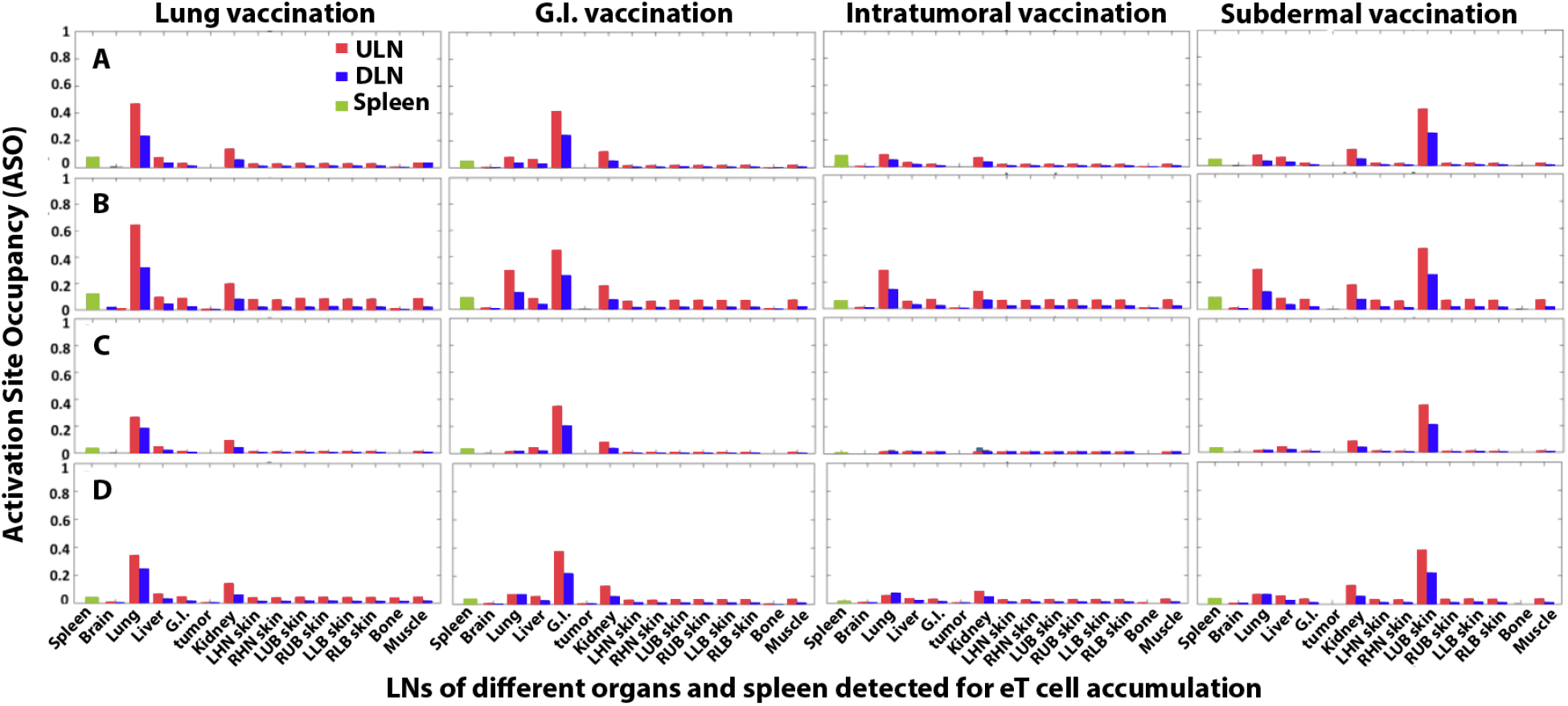
Accumulation of eT cells two weeks post immunization in the spleen and LNs of each organ defined as activation site occupancy (ASO). From left to right, the vaccination sites are lung, G.I., tumor, and left upper body (LUB) subdermal. A) Patients with low levels of tumor-induced antigen and suppressive factor (SF). B) Patients with 20% more tumor-induced antigen and the same level of SF as patients in A. C) Patients with the same level of tumor-induced antigen as patients in A but 20% increase in SF. D) Patients with 20% increase of both tumor-induced antigen and SF.

For all vaccination routes except intratumoral, ASO is highest in the LNs draining the vaccination site, and the difference between the vaccination LNs and other LNs increases either with higher SF levels (compare patient A with C in Fig.4) or with reduced tumor-produced antigen (compare patient B with A in Fig.4). In contrast, for intratumoral vaccination, the accumulation of SF in the tumor-draining LNs (Fig. 3) leads to early suppression of either APC activation or T cell activation by activated APCs and results in minimal ASO. In simulations with elevated tumor antigen production, both local and systemic T cell activation are enhanced, leading to higher ASO in vaccination site LNs, other LNs, and spleen (compare patient B with A in Fig.4). In contrast, elevated levels of tumor-derived SF significantly impairs T cell activation in the LNs near the vaccination site and severely reduce activation in most LNs and the spleen—particularly when vaccination occurred within the tumor, which is breast in these simulations (compare patient A with C in Fig. 4).

Comparing patients with the same rate of tumor antigen production, those with high SF levels have markedly different responses depending on tumor location and its connectivity to other organs and their respective LNs (patients A and B in Fig.4). For these patients, the farther the vaccination site from the tumor, the more robust the immune response, and intratumoral vaccination elicited a poor response. This is due to the accumulation of SF in tumor draining lymph nodes, which inhibits antigen-presenting cells (APCs) and directly blocks T cell activation.

As expected, increasing tumor antigen production when SF production is also high can mitigate the suppressive effects of the suppressive factor, restoring T cell activation levels to those observed for low SF production and low-antigen production (patients A and D in Fig.4). This suggests that patients with high antigen levels are less sensitive to immunosuppression and the location of the vaccination site, as the systemic release of tumor antigens can broadly activate LNs, with vaccination serving as only an additional antigen source—particularly in distal LNs, which are less affected by SF.

These findings suggest that, in patients with tumors producing a high level of suppressive factor, dual-site vaccination (both intratumoral (similar to when tumor releases antigen itself) and at a distant site) may improve T cell activation. Overall, the temporal dynamics and relative amounts of antigen and SF secretion by the tumor, and the relative anatomic location of the vaccine to the tumor site, dictate the sensitivity of the systemic immune response to vaccination. Therefore, simulating patient-specific responses based on tumor antigen and SF secretion rates may be necessary to optimize vaccination strategies.

While ASO indicates the best location(s) to apply the vaccine, OIH allows us to analyze immune responses based on vaccination site and timing, reflecting the body’s overall capacity to mount an anti-tumor response. Figure 5 presents OIH values two weeks after immunization (administered on day 7, 9, 11 or 13) for various internal organs and six superficial regions of skin across different patient profiles, categorized by the levels of tumor-induced antigen and SF. Results show that intradermal and intramuscular vaccinations produce similar OIH levels at each vaccination timing, regardless of site, indicating both are effective delivery routes. In high-SF patients, for early vaccination at day 7, the most effective internal vaccination sites (by descending OIH) are: G.I., spleen, bone, brain, lungs, tumor, kidney, and liver (patient C in Fig.5). Late vaccination attenuates this order and equalizes the responses from different vaccination sites. In low-SF patients (patients A and B)—particularly those with high antigen levels (patients B)—the lung yields the strongest response. These patterns highlight the advantage of immunization strategies based on tumor antigen and SF profiles. Immunization timing also influences vaccination outcomes: repeating simulations at different time points revealed distinct OIH dynamics, underscoring the value of personalized scheduling in cancer immunotherapy.

**Figure 5.**
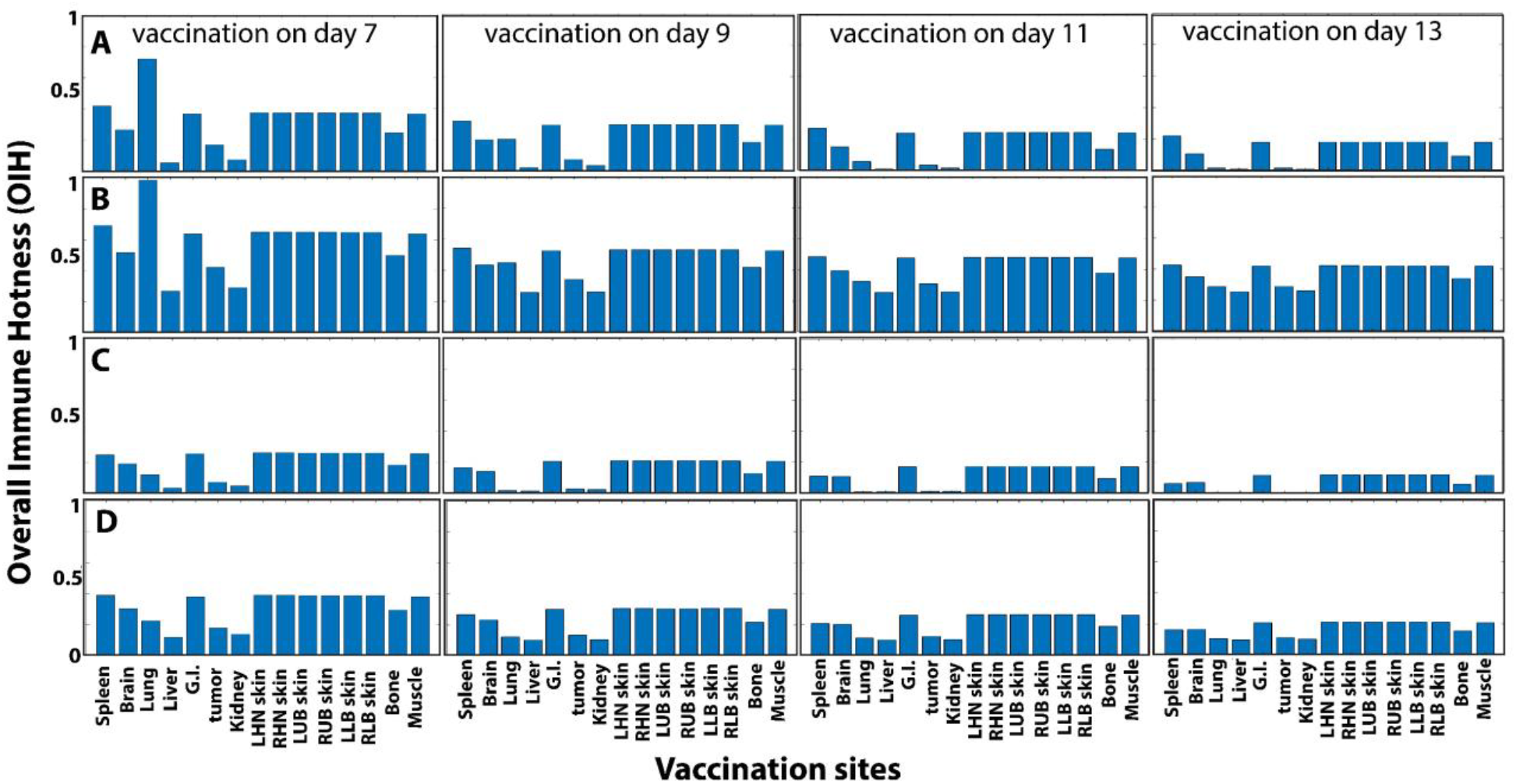
Overall Immune Hotness (OIH) for immunization in different sites at different times post tumor generation. A) Patients with low levels of tumor-induced antigen and suppressive factor (SF). B) Patients with 20% more tumor-induced antigen and the same level of SF as patients in A. C) Patients with the same level of tumor-induced antigen as patients in A but 20% increase in SF. D) Patients with 20% increase of both tumor-induced antigen and SF.

The ultimate goal of cancer vaccination is to increase the number and efficacy of anti-cancer immune cells in the tumor. To determine how vaccination affects eT cells in the tumor, we calculate the metric TH (Figure 6). The heat maps present TH as a function of vaccination site and timing for patients with variable tumor-induced antigen and SFs. TH is highly sensitive to the vaccination site during early stages of tumor development; however, this sensitivity progressively declines over time. At later stages, TH becomes largely independent of the vaccination site. The results show that even when vaccination is administered at an early stage of tumor progression, the SF secretion rate can significantly influence TH, effectively shifting the immune response from fully activated to completely suppressed. Furthermore, elevated SF levels diminish the sensitivity of T cell activation to both the timing and site of vaccination (Figure 6).

**Figure 6.**
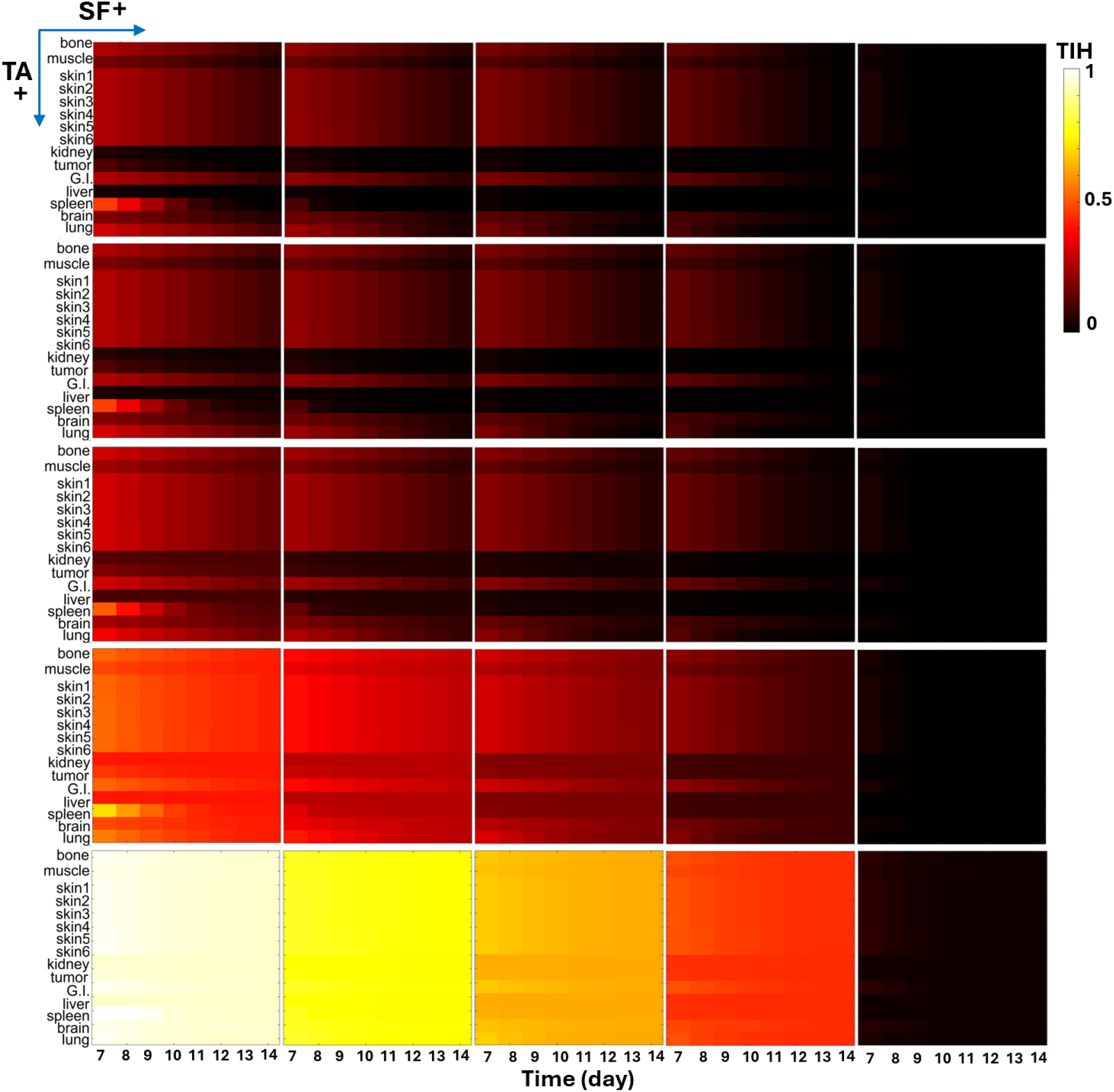
Tumor Immune Hotness (TIH) following immunization at different anatomical sites and time points, under varying levels of tumor-derived suppressive factors (SF) and tumor antigen (TA). From left to right, SF levels increase in 20% increments; from top to bottom, TA increases in 20% increments. Each cell in the grid represents the area under the curve (AUC) of normalized T cell counts in the tumor microenvironment (TME) during the two weeks following immunization at a specific site and time point. Skins 1 to 6 are respectively LHN, RHN, LUB, RUB, LLB, and RLB.

At low SF levels, early vaccination directly within the tumor can still elicit a measurable response, with spleen-targeted vaccination generating the strongest activation. However, as SF levels rise, the efficacy of spleen-based vaccination is markedly reduced, and the tumor becomes an ineffective site for inducing immune responses. This is because tumor-draining LNs (TDLNs) are the first lymph nodes to receive tumor-derived SF. However, TDLNs also contain high levels of tumor-associated antigens. Under conditions of low SF, intratumoral vaccination can further elevate antigen levels in the TDLNs, increasing the likelihood of APC and naïve T cell activation. In the spleen, where both antigen and naïve T cells are abundant, vaccination may accelerate T cell activation—provided that the high levels of tumor-derived SF have not already suppressed immunity.

Increasing tumor antigen levels at any given SF concentration enhances T cell activation and significantly reduces the dependence of immune activation on both vaccination site and timing. Nevertheless, high SF levels continue to suppress overall activation, even in the presence of abundant tumor antigen.

Given the wide range of conditions—including tumor antigen levels, suppressive factor concentrations, and variations in vaccination timing and site—simulating each new case or patient with the mechanistic model can be computationally intensive. To address this, we developed a reliable AI model that predicts outcomes for new patients based on prior simulation data. Specifically, we constructed a hybrid modeling framework that trains a random forest (RF) algorithm using our mechanistic PKPD model. The performance and predictive accuracy of this hybrid model are shown in Fig. 7A. The RF model demonstrated excellent agreement with the mechanistic model, with a validation coefficient of R = 0.99.

**Figure 7.**
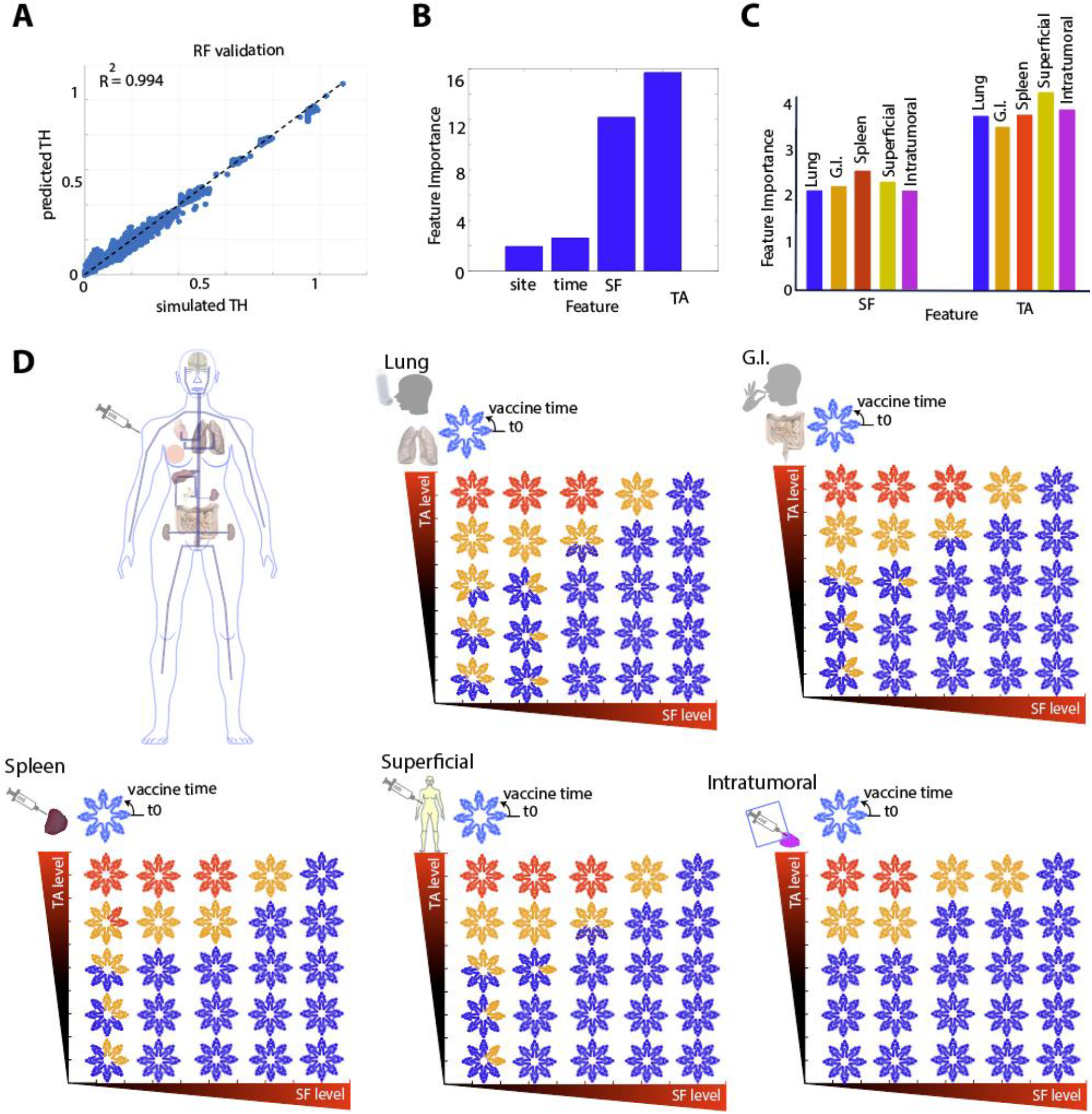
RF model and feature importance for vaccination outcomes. A) RF model developed from mechanistic model predictions of TH. B) Sensitivity of TH to vaccination site and timing under different SF and TA levels. C) Site-specific sensitivity of vaccinations to SF and TA for common sites including lung, G.I., superficial, and intratumoral, as well as spleen, which is unique in its potential for infusion and non-linear activation due to high immune cell and agent levels. D) Probability plots for vaccinations in lung, G.I., spleen, superficial, and intratumoral.

Using the trained RF model, we analyzed the sensitivity of TH to vaccination site and timing under varying levels of SF and TA, as shown in the feature importance plot in Fig. 7B. TA is the most important feature, followed by SF, then vaccination timing and site. For selective vaccination sites such as lung, G.I., spleen, superficial, and intratumoral, feature importance is shown in Fig. 7C, where TA is dominant across all vaccination routes, with the highest sensitivity for superficial vaccination and the lowest for G.I. In terms of SF, spleen vaccination is most sensitive, followed by superficial, G.I., lung, and finally intratumoral. Vaccination timing plays a reduced role in this analysis, allowing characterization of vaccination site effects for each patient with known TA and SF levels, especially when timing is uncertain due to unknown cancer stage. At the minimum, selecting the right vaccination site may enhance systemic immunity outcomes (TH).

As shown here, vaccination timing is a floating parameter dependent on cancer stage, making it a persistent challenge since timely tumor detection is often difficult. To address this, we aimed to estimate the probability of immune response to vaccination over time by repeating vaccination at multiple time points for each patient. In Fig. 7D, for each level of SF and TA, simulated patients were vaccinated on eight different days in the lung, G.I., spleen, superficial tissue, and tumor. Using k-means clustering, responses were grouped into three categories: non-responder (blue), semi-responder (orange), and responder (red). For known SF and TA levels, immunity probabilities are shown in Fig. 7D with cases corresponding to each SF and TA level indicated by different colors. For lung vaccination, the distribution was 12% responders, 23.5% semi-responders, and 64.5% non-responders. For G.I., 12% responders, 19% semi-responders, and 69% non-responders. For spleen, 13% responders, 20% semi-responders, and 67% non-responders. For superficial, 12% responders, 19% semi-responders, and 69% non-responders. For intratumoral, 8% responders, 16% semi-responders, and 76% non-responders. Assuming responder probability = 1, semi-responder = 0.5, and non-responder = 0, the overall vaccination efficacy rates for lung, G.I., spleen, superficial, and intratumoral sites were 23.75%, 21.75%, 23%, 21.5%, and 16%, respectively. Therefore, for unknown SF and TA, the suggested descending order of vaccination site efficacy is: lung, spleen, G.I./superficial, and lastly intratumoral.

## Conclusions

Tumor vaccinations hold considerable promise in clinical applications, yet their outcomes remain inconsistent. Critical factors for effective immune activation include: (a) the colocalization of antigen with immature APCs, and (b) the subsequent colocalization of antigen with activated APCs and naïve T cells, which typically occur in secondary lymphoid organs such as lymph nodes and the spleen. However, tumor-derived suppressive factors can disrupt these activation processes and dampen anti-tumor immunity. Our findings demonstrate that the spatial distributions of antigen, immune cells, and suppressive factors strongly influence both the timing and location of T-cell activation, as well as their eventual accumulation within the tumor. While tumor-driven production of antigen and suppressive mediators plays a central role in initiating immune responses, carefully optimizing the timing and spatial delivery of vaccines may substantially enhance the efficacy of anti-tumor vaccination strategies.

The computational framework developed in this study represents a first-of-its-kind multiscale PBPK model that integrates comprehensive systemic circulation with coupled superficial–inner organ networks and a sequential lymph node architecture capturing upstream and downstream lymphatic drainage—features essential for localized immune activation in distinct lymphosomes. The model mechanistically tracks antigen, suppressive factors, naïve and effector T cells, and immature and activated APCs, along with their phenotype-dependent cytokine release, across both vascular and lymphatic pathways. It also incorporates organ-specific infiltration dynamics through mechanistic fluxes. Developed using a dynamic matrix-based approach, the framework can be readily adapted to other species, such as mouse, facilitating translational studies that bridge preclinical and human systems. Unlike conventional PBPK models validated with unrelated species data, this model was verified through a multi-step, human-relevant validation process spanning cellular interactions to whole-body immune responses.

Overall, this mechanistic and human-validated framework provides quantitative, predictive insights into the competition between immune activation and tumor-derived suppression, establishing a Digital Twin platform to simulate and optimize personalized vaccination routes, schedules, and timing for improved cancer immunotherapy outcomes.

## Acknowledgements

Lance Munn’s research is supported by NIH R01CA247441, U01CA261842 and R21EB031982. Timothy Padera’s research is supported by NIH R01CA284372, and R21AG072205. Munn and Padera’s research is supported by NIH R01CA284603, and R01HL128168.

